# Action Imagery and Observation in Neurorehabilitation for Parkinson’s Disease (ACTION-PD): development and pilot randomised controlled trial of a user-informed home training intervention to improve everyday functional actions

**DOI:** 10.1101/2020.07.14.188375

**Authors:** Judith Bek, Paul S. Holmes, Chesney E. Craig, Zoë C. Franklin, Matthew Sullivan, Jordan Webb, Trevor J. Crawford, Stefan Vogt, Emma Gowen, Ellen Poliakoff

## Abstract

**Background:** Parkinson’s disease (PD) causes difficulties with everyday manual activities, but few studies have addressed these therapeutically. Training with action observation (AO) and motor imagery (MI) can significantly improve performance in healthy individuals, particularly when these techniques are applied simultaneously (AO+MI). Both AO and MI have shown promising effects in PD, but previous studies have used these separately. This article describes the development and pilot testing of an intervention combining AO+MI and physical practice to improve functional manual actions in PD.

**Methods:** The home-based intervention, delivered using a tablet computer app, was iteratively designed by an interdisciplinary team including people with PD, and further developed through focus groups and initial testing. The intervention was then tested in a six-week randomised controlled trial (ISRCTN 11184024) of 10 participants with mild to moderate PD (6 intervention; 4 treatment as usual).

**Results and Conclusions:** Usage and qualitative data provided preliminary evidence of acceptability and usability, indicating that a feasibility RCT is warranted. Exploratory analyses suggested potential improvements in manual actions. The importance of personalisation, choice, and motivation was highlighted, as well as the need to facilitate engagement in motor imagery. The findings also have broader relevance for AO+MI interventions in PD and other populations.

## Introduction

Beyond the more widely recognised difficulties with gait, balance and gross motor functioning, Parkinson’s disease (PD) impairs fine motor skills including hand dexterity, which are needed for the successful performance of activities of daily living [e.g., 1,2]. Sudden arrests in movement – known as “freezing” – of the upper limbs can also occur in PD, which may be correlated with freezing of gait [3]. Daily activities can be affected even in the early stages of PD [4], potentially impacting on work performance as well as household tasks, self-care, hobbies and leisure activities; thereby significantly limiting independence and quality of life. Indeed, people with PD consistently report dexterity among the domains most affected by the condition [5,6], and have expressed a need for interventions to improve dexterity [7,8]. However, few studies have directly addressed dexterity problems in PD [9]. One large trial of home-based dexterity training for people with PD used task-specific hand exercises with written and illustrated instructions [10]. Short-term improvements in dexterity and activities of daily living were found when compared with a control group undergoing resistance training, but these effects were not sustained at 12 week follow-up.

One general approach to facilitating movement in people with PD is the use of external cues such as visual stimuli (e.g., floor markers or laser pointers) and auditory stimuli (e.g., metronome beats or rhythmic music). Although PD affects the internal generation of action [11], external cues can help to elicit or control movement; this may relate to the relative preservation of goal-directed movement pathways, which compensate for impaired habitual or automatic processes [12]. External sensory cues are widely used by therapists to improve walking speed or other gait characteristics in PD, and are supported by empirical evidence [13,14]. However, these cues are less applicable to the fine hand movements required for everyday functional actions. Additionally, they cannot always be readily applied in real-life situations outside of the clinic or laboratory, and long-term effects of cueing have not been established [14].

An alternative type of movement cue may be provided by observation of human action (action observation; AO). A large body of literature based on investigations in healthy participants has demonstrated that AO facilitates movement and increases motor learning [15–18]. This involves the activation of an action observation network [19], incorporating a set of fronto-parietal neural structures that are engaged during both AO and motor execution, referred to as the “mirror neuron” system. Motor imagery (MI) is another process that shares neural substrates with AO and motor execution [20]. MI is the imagination of movement in the absence of overt action [21], with associated sensations (kinaesthetic imagery) and images (visual imagery), which also facilitates learning and movement [22–24].

Until recently, research on AO and on MI has been undertaken by different scientific communities, and applications in sports and neurorehabilitation have focused on either AO or MI. However, Vogt and colleagues [24] suggested that individuals can engage in both forms of motor simulation simultaneously, by performing MI of an observed action. In healthy participants, this combined “AO+MI” has been found to produce greater behavioural and neurophysiological effects than either process in isolation [24–26] and preliminary evidence suggests that combined AO+MI may be effective in stroke rehabilitation [27]. Kinaesthetic imagery (focusing on sensation and effort) is typically emphasised in AO+MI interventions, and is associated with stronger sensorimotor activations than visual imagery [25].

Below, we introduce the rationale and evidence for using AO and MI in PD, and the background for an intervention utilising combined AO+ MI.

### Action observation and motor imagery as tools to facilitate movement in PD

AO and MI have been studied in the context of neurorehabilitation, primarily with stroke patients, with promising effects reported [28–30]. Moreover, MI training is recommended in rehabilitation guidelines from the American Stroke Association [31]. Although a smaller number of studies have investigated AO and MI in PD, AO has been found to influence movement speed and timing in reaching [32] and finger-tapping [33] tasks, as well as hand movement amplitude [34], and people with PD have shown preserved motor resonance for incidentally-observed hand actions [35]. People with PD also report similar vividness of MI to healthy controls; however, like their actual movements their imagery may be slowed [36], and compensatory mechanisms may be involved, such as a greater reliance on visual processes [37,38]. Additionally, evidence from spontaneous gestures when describing actions suggests that people with PD may rely more on the third-person perspective to internally represent movement [39]. This indicates that motor simulation may be reduced or more effortful [37,40].

Small-scale intervention studies in PD have provided preliminary evidence that AO combined with physical practice can improve motor symptoms, balance and gait [41,42], as well as dexterity [43] and functional independence [44]. Increased activation in cortical motor areas has also been found following AO-based training in PD [41], suggesting potential neuroplastic effects. MI has been found to help overcome freezing of gait in people with PD [45], and MI training combined with physical practice improved timed motor performance [46]. These findings suggest that both AO and MI may enhance or facilitate access to motor representations in people with PD.

Despite the evidence supporting combined AO+MI approaches in healthy participants and in stroke rehabilitation as described above, only one study to date has investigated AO+MI in PD, showing increased imitation of hand movements when participants engaged in MI during AO, compared to AO alone [34]. It has been proposed that combining AO and MI may increase corticospinal excitability in people with PD, thereby enhancing pre-movement facilitation [47]. Additionally, concurrent observation provides an ongoing visual input, which may facilitate the generation of motor imagery (see [25]), potentially compensating for difficulties with MI in people with PD [34]. Indeed, spontaneous MI may have contributed to the effects of previous AO interventions that did not include explicit MI instructions.

Visual perspective is also an important consideration in AO and MI interventions. In particular, observation of actions from the first-person perspective (as if watching through the actor’s eyes) is thought to promote kinaesthetic imagery (e.g., [48]), compared with the third-person perspective (watching the actor from an external position). Accordingly, in neuroimaging studies of healthy individuals, first-person imitation has been associated with increased activation of the sensorimotor cortex [49] and recruitment of fewer neural regions [50], suggesting reduced difficulty compared with the third-person perspective. However, the third-person perspective may provide useful information about the context and overall movement parameters of an action (e.g., [51]). Previous AO intervention studies in PD have shown positive effects using both first-person videos [43] and third-person videos [33,41,42,52].

### A user-informed, personalised home-based AO+MI intervention for PD

To investigate the potential of combined AO+MI training to improve everyday activities in mild to moderate PD, we designed the ACTION-PD intervention, which utilises video-based AO+MI and physical practice of functional manual actions, delivered via an app on a tablet computer. To ensure that any intervention is relevant, feasible and acceptable in the target population, it is important to consult with potential users [53], and to conduct qualitative research to inform development, as recommended by the UK Medical Research Council (MRC) guidelines for complex interventions [54]. We involved people with PD in the development of the intervention through focus groups and as members of the research team.

Given the heterogeneous nature of PD, “personalised treatments” has been identified as a research priority by people with PD [8] to address the broad range of everyday activities that individuals may find difficult (e.g., [7]). In this respect, training based on action representation (AO and MI) can be tailored to the individual’s needs and rehabilitation goals. While the ultimate aim of the intervention is to develop skills in using AO+MI that individuals can apply across multiple situations, focusing on personally meaningful actions is likely to increase motivation and engagement with the training [7].

ACTION-PD also differs from other interventions by providing a home-based option, whereas in previous studies, AO interventions were conducted in clinics or under physiotherapist supervision (e.g., [41–43]). The feasibility of other home interventions using digital technology in PD has been reported, such as “exergame” activities focused on movement timing and coordination (e.g., [55,56]), and if effective, this approach could provide a widely accessible, low-cost alternative or supplement to existing rehabilitation programmes.

A previous focus group with people with PD [7] explored views on the proposed ACTION-PD intervention, as well as individuals’ understanding and experiences of AO and MI. The focus group indicated that a home-based combined AO+MI intervention would be acceptable and useful for people with mild to moderate PD. This study also reiterated the importance of personalisation and flexibility, as well as highlighting the need to consider the issue of motivation.

In this article we describe the next stages in the development and pilot testing of the intervention, which consisted of: (i) design of the intervention prototype; (ii) a further focus group to obtain feedback on the prototype and to explore potential barriers to technology use among people with PD; (iii) initial field testing; and (iv) a pilot randomised controlled trial (RCT). Our aim was to collect preliminary qualitative and quantitative data on usability, acceptability and potential outcomes of the intervention, in order to establish whether a feasibility RCT is warranted. The intervention development process from conceptualisation to pilot testing is outlined in Figure 1.

**Figure 1.**
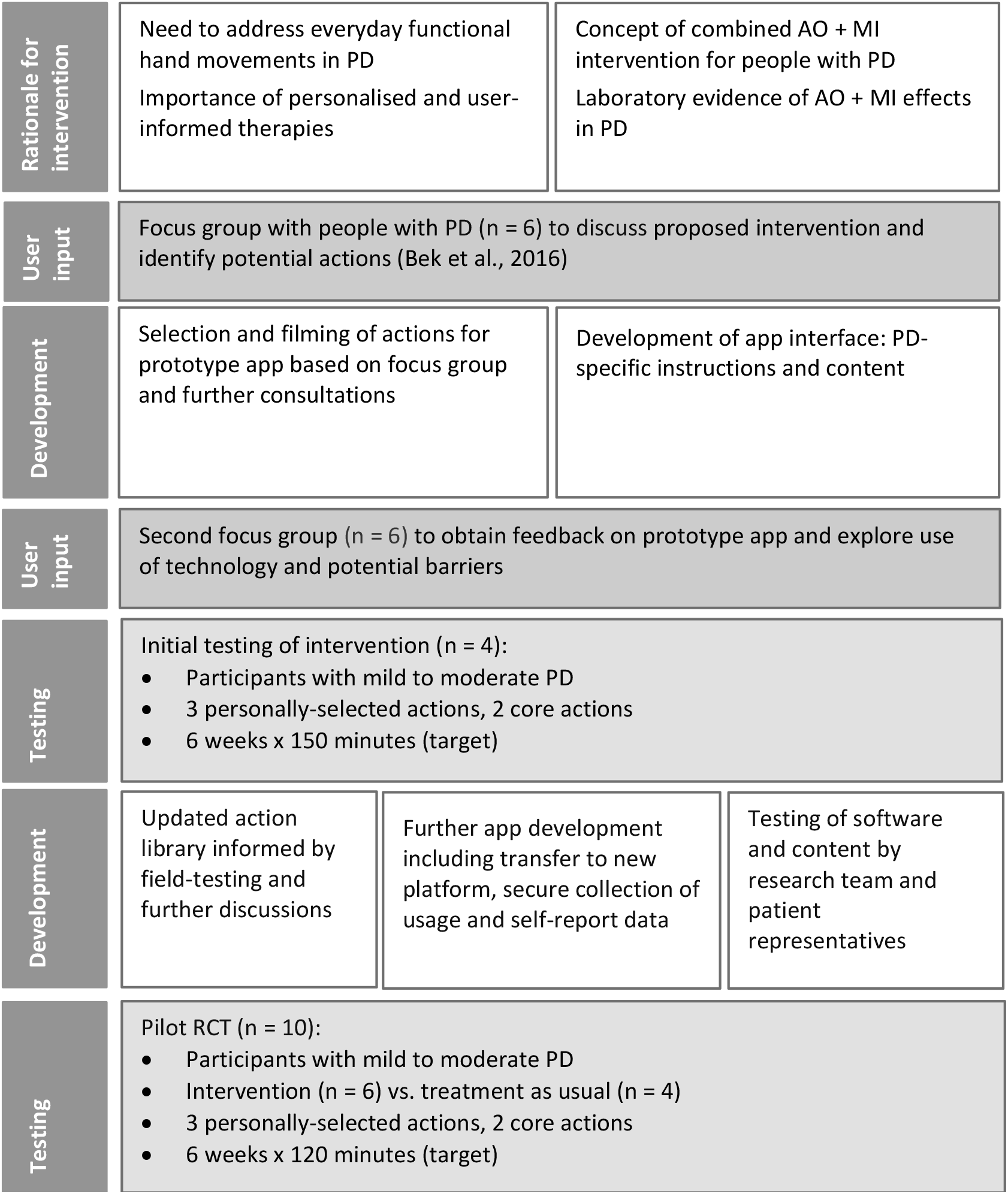
The intervention development process.

## Intervention development

### The intervention prototype

An action library was first compiled to enable users to select the actions they wished to train: actions were identified from suggestions provided in the previous focus group [7], as well as examination of the literature and discussions within the research team. The selection of actions was limited to those that could be practiced safely at home in a seated position, using everyday objects. Patient representatives were invited to provide feedback on the initial selection, and to suggest any additional actions.

The actions selected to include in the prototype (see Figure 2 for examples) were video-recorded in a quiet room, using a plain wooden table and a neutral background free from other objects or distracting features.

**Figure 2.**
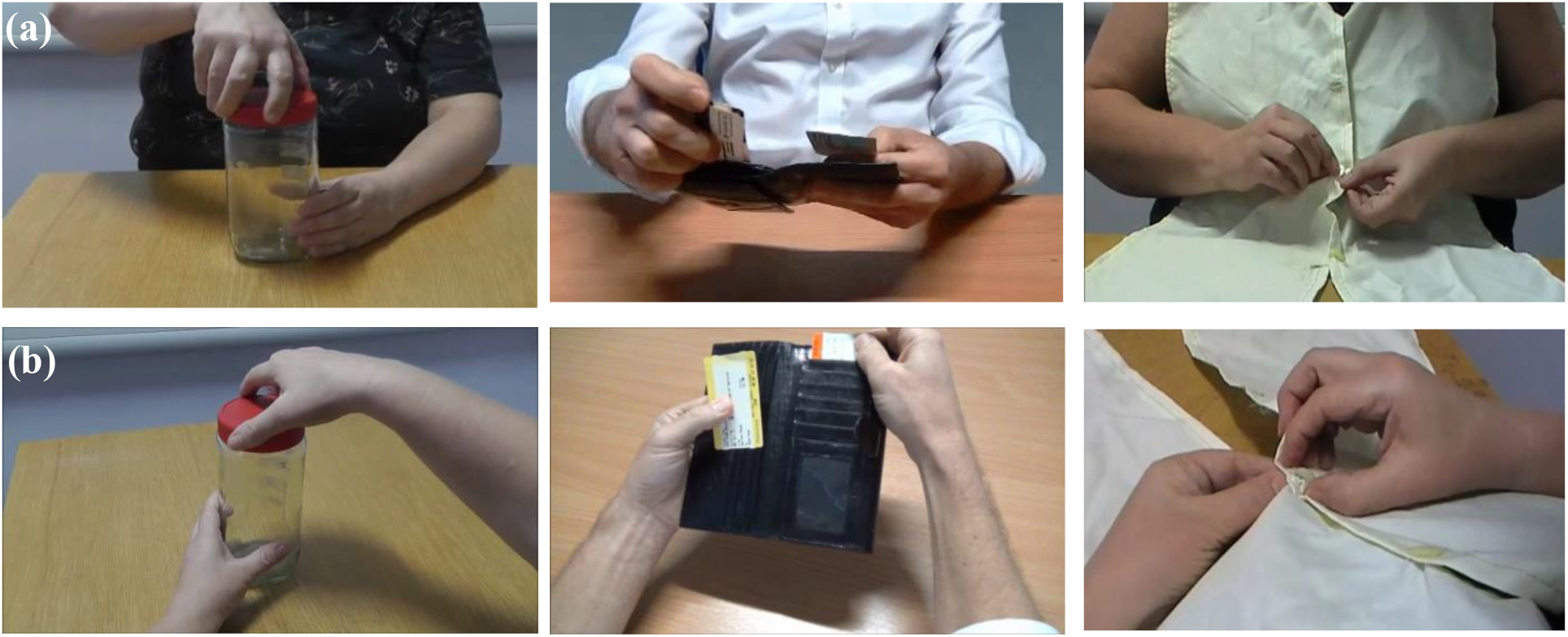
Examples of everyday actions used in the intervention (coffee jar, ticket sorting, buttoning). Each action is presented from the third-person perspective (a) followed by the first-person perspective (b).

Each action was filmed with male and female actors to allow matching to the participant’s gender, and from both third-person and first-person perspectives. The third-person video was filmed from either the front or side of the actor, depending on which perspective provided the clearest view of the action. This provided the overall context of the action and movement kinematics, while the first-person perspective was expected to promote kinaesthetic imagery and enhance sensorimotor activations (e.g., [49]).

The prototype was developed through modification of an app originally designed for upper limb rehabilitation in stroke patients based on observing, imagining and then executing actions [57]. For ACTION-PD, the app was updated with the PD-relevant videos, as well as instructions for simultaneous rather than sequential AO and MI. The third-person video of the action was presented first, followed immediately by the first-person video (see Figure 3). Videos were played with the accompanying sound, which provides additional action-relevant information, and may evoke auditory activation of sensorimotor areas and facilitate motor imagery (e.g., [58,59]). Participants were instructed to observe the videos while simultaneously engaging in motor imagery, followed immediately by physical execution of the action using the same objects as depicted in the video. During action execution, a still image of the action (extracted from the first-person video) was displayed on the screen as a reminder. This remained on screen for the same duration as the preceding video, but participants were advised that they were not required to complete the action within this time limit.

**Figure 3.**
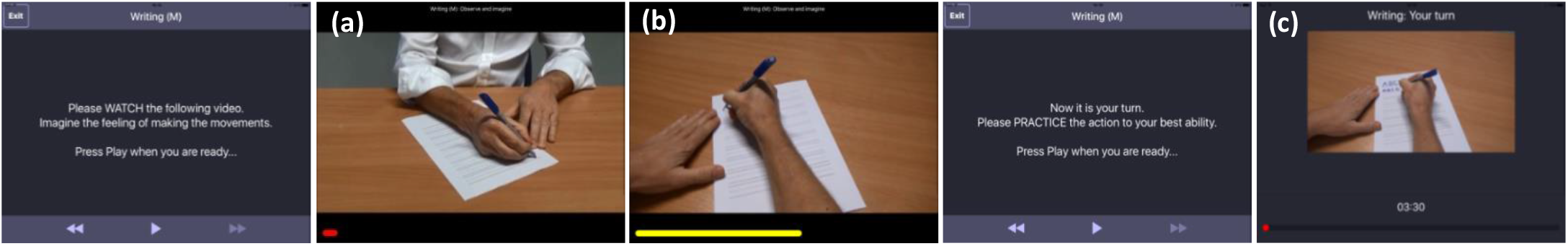
Screenshots of the prototype app used in the pilot RCT: Participants were instructed to imagine each action (kinaesthetic motor imagery) while watching videos showing the action from the third-person (a) and first-person (b) perspectives, before physically performing the action using the relevant objects (e.g. pen and paper). A still image of the action (c) was displayed during action execution. Finally, participants rated the vividness of their imagery during observation and the difficulty of performing the action.

### Focus group

To obtain feedback on the intervention prototype and explore views and experiences of technology, a focus group was conducted with 6 participants with mild to moderate PD (Hoehn & Yahr stage 1 to 3; see Table 1 for demographic characteristics). Five of the participants had attended our previous focus group [7] and one was a member of the research team; all participants were therefore already familiar with the concepts of AO and MI and the principles underlying the intervention.

**Table 1:** Demographics of focus group participants.

| Participant | Sex | Age<br>(years) | Time since diagnosis<br>(years) |
| --- | --- | --- | --- |
| FG1 | M | 53 | 7 |
| FG2 | M | 61 | 21 |
| FG3 | M | 70 | 7 |
| FG4 | F | 57 | 9.5 |
| FG5 | F | 64 | 3.8 |
| FG6 | M | 65 | 3.2 |

The focus group was chaired by one of the authors (EP) and facilitated by two others (JB and JW). A schedule of topics was used to guide discussions, with open-ended questions and additional prompts where needed. The prototype app was demonstrated and participants were then invited to try the app themselves and comment on the interface and functionality. The proposed training protocol was also discussed, as well as the use of technology more broadly, including participants’ experience in using mobile devices and apps, and potential symptomatic barriers to technology use. Responses were recorded and transcribed by an independent transcription service.

Using thematic analysis with an inductive approach [60], themes relating to (i) accessibility, (ii) motivation, and (iii) personalisation and flexibility, were identified. The findings are summarised in Table 2; a full list of themes with illustrative quotes is provided in the supplementary material (S1).

**Table 2:**
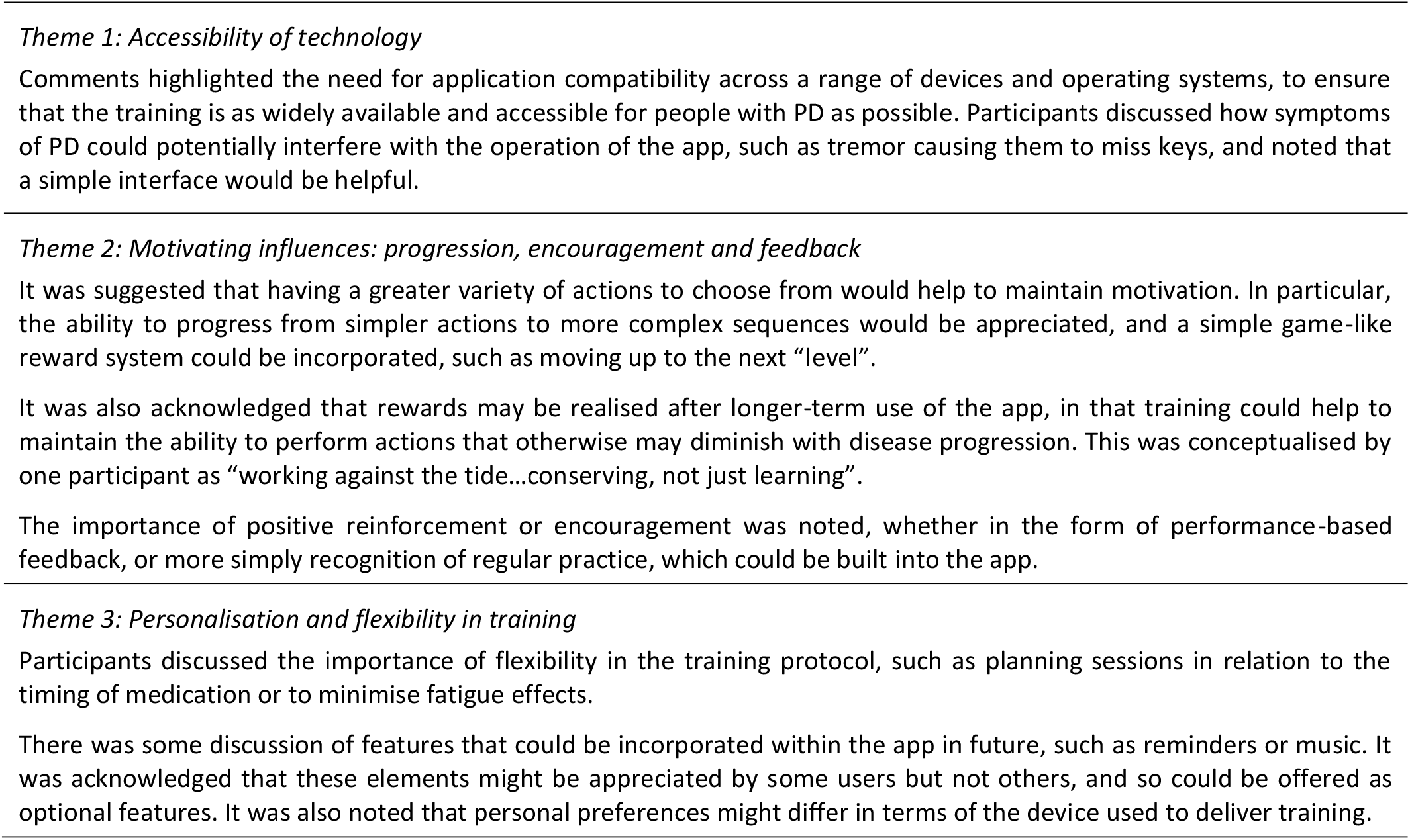
Themes identified from the focus group.

## Initial testing and pilot RCT

Following the overall positive feedback from the focus group, the prototype intervention was pilot-tested to explore usability and acceptability, as well as potential outcomes in terms of dexterity, reaction times, motor imagery and quality of life. It should be noted that due to software and time constraints, it was not possible to implement all the suggestions from the focus group at this stage, including progression of actions and motivating elements. Testing was conducted in two stages: (i) initial testing with a small number of participants; (ii) a pilot RCT. Below we report the methods and results of both stages together, indicating where changes were made between the initial testing and pilot RCT.

### Participants

For the initial testing phase, four participants with mild to moderate PD and with no history of other neurological or psychiatric conditions were recruited from a volunteer panel and through Parkinson’s UK (see Table 3). Participants reported experiencing difficulties with everyday manual actions, had normal or corrected-to-normal vision and were screened for cognitive impairment using the Addenbrookes Cognitive Examination (ACE-III [61]). For the pilot RCT, a further 10 participants with mild to moderate PD were recruited and screened in the same way (Table 3).

**Table 3.** Baseline characteristics of participants in pilot testing.

| Participant | Sex | Age (years) | Time since diagnosis (years) | Hoehn & Yahr stage | UPDRS-III motor score |
| --- | --- | --- | --- | --- | --- |
| Initial_1 | M | 73 | 7 | 2 | 54 |
| Initial_2 | F | 72 | 10 | 3 | 36 |
| Initial_3 | M | 63 | 8 | 1 | 16 |
| Initial_4 | F | 50 | 2 | 2 | 32 |
| RCT_I1 | M | 70 | 4 | 2 | 49 |
| RCT_I2 | M | 65 | 7 | 2 | 29 |
| RCT_I3 | M | 71 | 4 | 2 | 40 |
| RCT_I4 | M | 66 | 16 | 2 | 37 |
| RCT_I5 | F | 69 | 2 | 3 | 47 |
| RCT_I6 | M | 60 | 2 | 3 | 66 |
| RCT_C1 | M | 66 | 13 | 2 | 51 |
| RCT_C2 | M | 59 | 5 | 2 | 39 |
| RCT_C3 | M | 63 | 2 | 1 | 28 |
| RCT_C4 | M | 47 | 4 | 2 | 42 |

Note: Initial = initial testing cohort; RCT\_I = pilot RCT intervention group; RCT\_C = pilot RCT control group.

### Design and protocol

#### Initial testing

With the assistance of a researcher, each participant selected 3 “personal” actions they wished to improve (e.g., buttoning, writing, opening and closing food containers). In addition, to explore the possibility of a more standardised approach to training and outcome measurement, all participants were asked to practice two “core” actions selected by the research team, which were based on common everyday tasks (handling coins, sorting train tickets). The combined videos (first- and third-person perspectives) had a mean duration of 54.9 s. A full list of personal and core actions is provided in the supplementary material (S2).

Following a baseline assessment in the laboratory (see “Outcome measures” below), a researcher visited the participant at home to deliver the tablet computer and accessory objects corresponding to the items used in the videos, and to demonstrate the use of the app and explain the training protocol. A full instruction guide, as well as background information on the project and contact details for the research team, was provided within the app. Participants were also given a printed copy of the instructions. The researcher answered any questions and ensured that the participant fully understood how to use the app before independent training commenced.

The training was carried out in the individual’s home for 6 weeks using the app on a tablet computer (iPad). In each training session, participants practiced the 5 actions (3 personal and 2 core), which were presented in a randomised order to avoid fatigue disproportionately affecting performance or completion of some of the actions. A target training time of 150 minutes per week was set (based on previous action observation intervention studies; [30]), which could be divided up according to the individual’s preference. For example, if a single training session took 25 minutes, the participant could choose to complete one session per day for 6 days, or two sessions per day for 3 days. To maximise feasibility, the training was intended to be flexible, and participants were advised that they could fit their practice around other commitments or difficulties relating to symptoms.

Participants were asked to record dates and times of practice sessions in a paper-based training diary. For each session, they were also asked to rate the difficulty of performing each action on a five-point scale (very easy/quite easy/neither easy nor difficult/quite difficult/very difficult). During the training period, participants were followed up with a weekly telephone call, and were also encouraged to contact the research team at any other time if they had questions or experienced technical issues.

On completion of the 6-week training period, participants returned to the laboratory for a follow-up assessment (approximately 10 weeks after baseline). Semi-structured interviews were then conducted to obtain qualitative feedback on the app and explore individuals’ experiences of the training.

#### Pilot RCT

The pilot RCT was registered with ISRCTN (trial number 11184024). The flow of participants through the trial is illustrated in a CONSORT diagram [62] in Figure 4. Prior to the pilot RCT, the app was transferred to a new software platform that enabled secure in-app collection and storage of usage and self-report data, in place of the paper-based training diaries used in the initial testing phase. A larger library of videos was also produced, based on feedback from the initial testing and further discussion within the research team. Additionally, two new “core” actions (opening and pouring from a water bottle, transferring sugar from a jar to a cup) were identified in discussion with Parkinson’s representatives.

**Figure 4.**
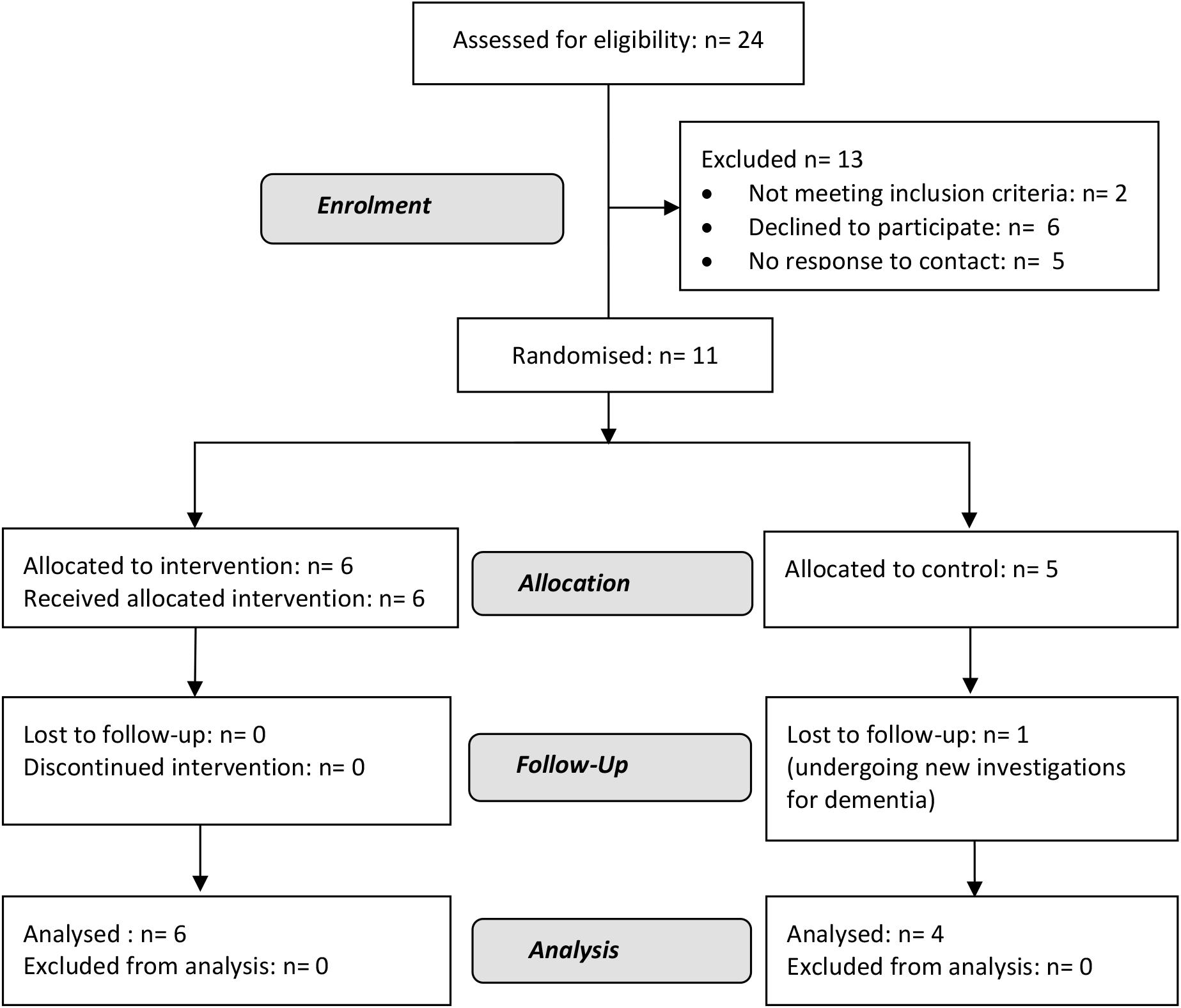
CONSORT diagram showing flow of participants in the pilot RCT.

Each participant selected six actions from the updated action library in order of preference: the first three actions were included in the individual’s training programme (“personal-trained”) while the other three (“personal-untrained”) were used to test for transfer of training effects. The two core actions were included in training for all participants.

Following baseline assessment, participants were randomly allocated to the intervention group or control group by a researcher who was not involved in recruitment or data collection, using an online randomisation tool. The intervention protocol was the same as described above except for the following:

i. Based on data from the initial testing suggesting that training sessions took less time than anticipated to complete, and that participants were not all achieving the weekly target, the training time was reduced to 120 minutes per week. Again, this could be divided up according to the participant’s preference (e.g., two 20-minute sessions per day for 3 days per week).
ii. Immediately after completing each action, participants were asked within the app to rate the vividness of their imagery when watching the video, using a five-point scale. The difficulty of the action was then also rated on a five-point scale.

The control group continued with their usual treatment for PD and did not receive the intervention, but were followed up with a weekly telephone call to maintain contact.

### Outcome measures

Usage data and action ratings were collected via the home training diaries (initial testing) or within the app (pilot RCT). In the pilot RCT, imagery ratings for each action were also collected within the app. Additional information on usability and acceptability was obtained through semi-structured post-training interviews, in which participants were asked about their experiences of the app and the training content and schedule, as well as any perceived changes in their performance of the actions and transfer of skills to other tasks. Where possible, baseline and follow-up assessments were conducted at the same time of day to minimise variability in relation to medication effects.

The following exploratory measures were included:

i. Dexterity was assessed using the Dexterity Questionnaire (DextQ-24 [63]); a self-report questionnaire designed for people with PD, which examines manual dexterity for a range of everyday tasks.
ii. Quality of life was assessed using the Parkinson’s Disease Questionnaire (PDQ-39 [64]).
iii. Motor imagery was tested using the Kinaesthetic and Visual Imagery Questionnaire (KVIQ; [65]), which has been validated in people with PD [66]. The KVIQ requires participants to physically perform, and then imagine performing, simple actions involving different body parts (upper limbs, lower limbs, trunk, shoulders and head). Visual and kinaesthetic subscales are used to rate the vividness of images and intensity of sensations respectively, each on a five-point scale.
iv. Simple and choice reaction time tests required participants to react to the appearance of an LED by pressing a button on a response box as quickly as possible (see [67] for details). The simple task consisted of two blocks, with responses made using the left hand in the first block and the right hand in the second. In the choice RT task participants responded using the hand corresponding to the location of the light signal, which appeared in a random order on either the left or right side of the display.

In the pilot RCT, performance of personalised (trained and untrained) and core actions was also assessed in the laboratory. Participants viewed videos showing each action from the third-person and then first-person perspective, while engaging in kinaesthetic imagery, before physically performing the action. Each action was presented 3 times, resulting in a total of 24 trials. Videos were viewed on a projector screen (300 × 580 mm display size), approximately 1100 mm from the participant, who was seated at a table with the objects needed to complete the action placed in front of them. The objects were occluded by an opaque screen until the end of the video, when a go-signal indicated the start of the physical practice as the objects were revealed (the word “Go” in text appeared on the screen, accompanied by a beep). Following each trial, participants were asked to rate the difficulty of performing the action on a five-point scale. Action performance was filmed using a video camera positioned adjacent to the projector screen, and the time taken to complete each action was coded from the video by a researcher who was blinded to group allocation.

## Results

### Feasibility

#### Training adherence

All participants in the initial testing and those in the intervention arm of the pilot RCT completed the 6 weeks of training, with an average of 7.8 (range: 5.7 - 11.7) sessions per week in the initial phase and 8.9 (6 – 14) sessions per week in the pilot RCT. Based on an estimated average session duration of 20 minutes, this corresponds to a mean adherence of 104 % in the initial cohort (76 – 156 %) and 148.3 % in the pilot RCT (99.5 – 233 %).

#### Post-training interviews

The semi-structured interviews were analysed thematically using the same approach as described above for the focus group. Given the overlap in content of the interviews, data from the initial testing phase and the pilot RCT were combined for analysis. Themes are summarised in Table 4 and a more detailed analysis with illustrative quotes is provided in the supplementary materials (S3). Following the interview, each participant was asked to rate aspects of the app and training on five-point scales. All participants rated the app usability and the actions as either “very easy” or “quite easy”, and said that they would “definitely” or “probably” use a similar app in the future. Eight of the ten participants reported that they enjoyed the training “very much” or “somewhat”, five felt that they had “definitely” or “probably” improved on the trained actions, and six reported that they had “probably” improved on other untrained actions.

**Table 4.**
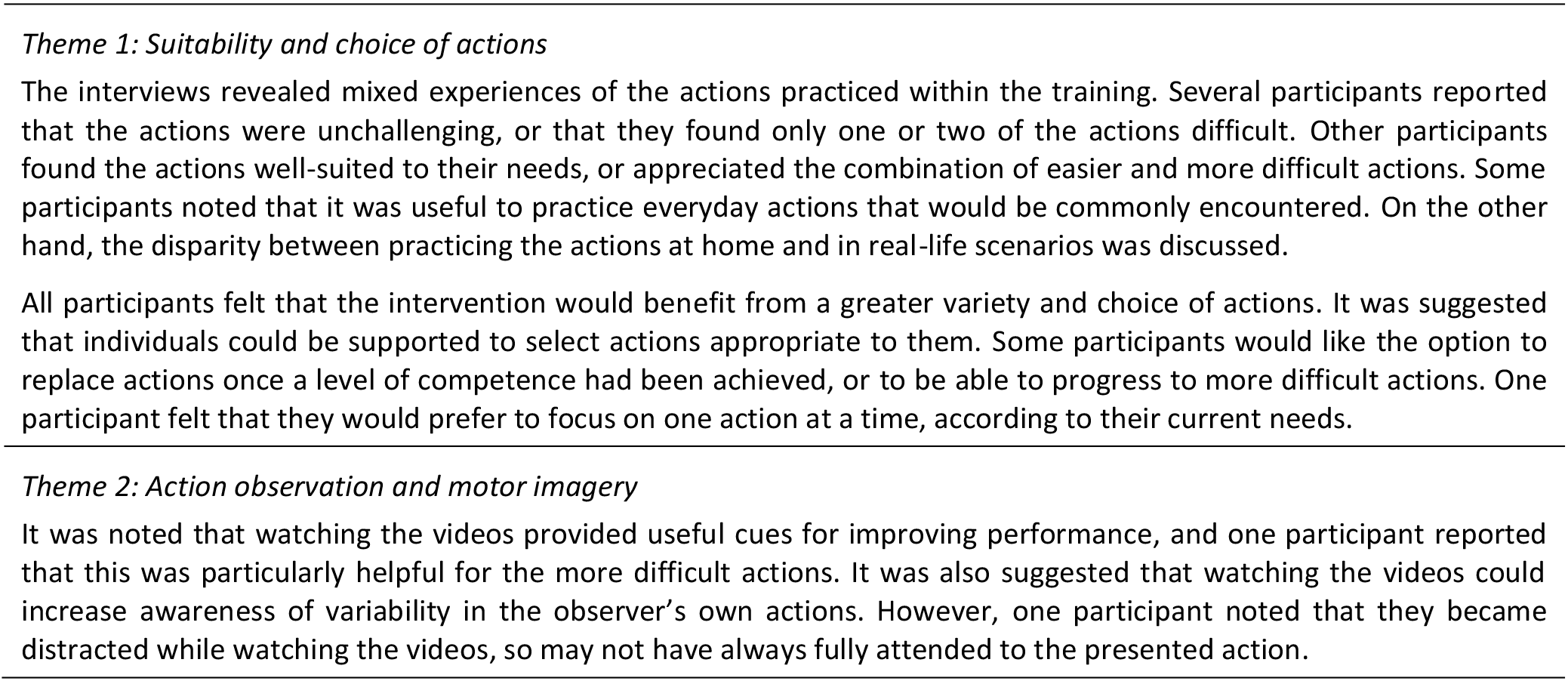
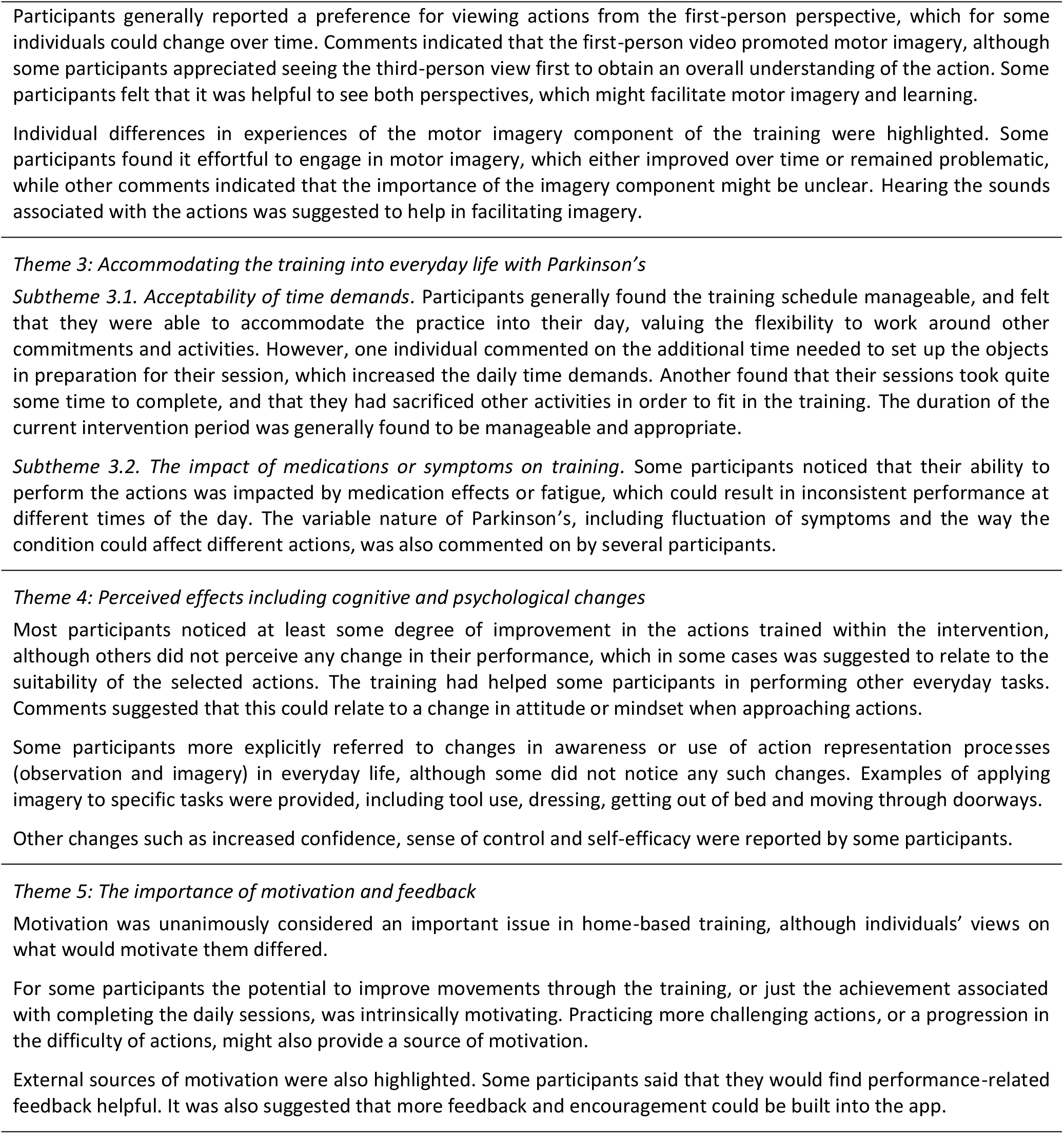
Themes generated from semi-structured post-training interviews.

### Action difficulty and motor imagery ratings

Ratings of action difficulty and motor imagery vividness during training are summarised in the supplementary materials (S4). Across the initial testing and pilot RCT, an overall reduction in difficulty ratings between the first and sixth weeks was found for both core actions (median change = 35.1 %) and personal actions (median change = 43.4 %). Core actions were rated as easier than personal actions from the start of training and perceived improvements appeared to reach a plateau by week 2 in both cohorts. In the pilot RCT, ratings of motor imagery did not show any evidence of improvement across the 6 weeks; in fact there was a slight reduction in reported vividness (median change = 16.2 %).

### Exploratory outcomes

Results of the baseline and follow-up laboratory assessments are presented in Table 5. Statistical analyses were not performed because of the small sample sizes. However, numerical data suggested some improvement in self-reported dexterity as well as simple and choice reaction times (see Figure 5).

**Table 5.** Performance on exploratory outcome measures in the initial testing and pilot RCT: median pre- and post-scores, change (baseline minus follow-up) and interquartile range of change score.

| Group | DextQ-24 | KVIQ | Simple RT (ms) | Choice RT (ms) | PDQ-39 |
| --- | --- | --- | --- | --- | --- |
| Initial testing | 36.5; 33.0<br>(2.0; 1.25) | 107.5; 105.5<br>(-7.50; 27.6) | 323; 296<br>(17.0; 42.0) | 409; 388<br>(-14.13; 83.19) | 22.50; 21.0<br>(-1.0; 6.75) |
| Pilot RCT: |  |  |  |  |  |
| Intervention | 36.5; 34.5<br>(2.5; 6.25) | 122.0; 117.0<br>(4.50; 22.8) | 309; 312<br>(-1.13; 34.13) | 428; 395<br>(18.25; 129.44) | 41.5; 36.0<br>(-1.50; 11.5) |
| Control | 26.0; 26.0<br>(0.0; 3.0) | 130.0; 118.0<br>(7.0; 30.8) | 356; 349<br>(-6.75; 34.13) | 395; 427<br>(-44.75; 52.13) | 13.50; 7.0<br>(2.50; 5.0) |

Note: Positive change values indicate improvement, except on the KVIQ.

**Figure 5.**
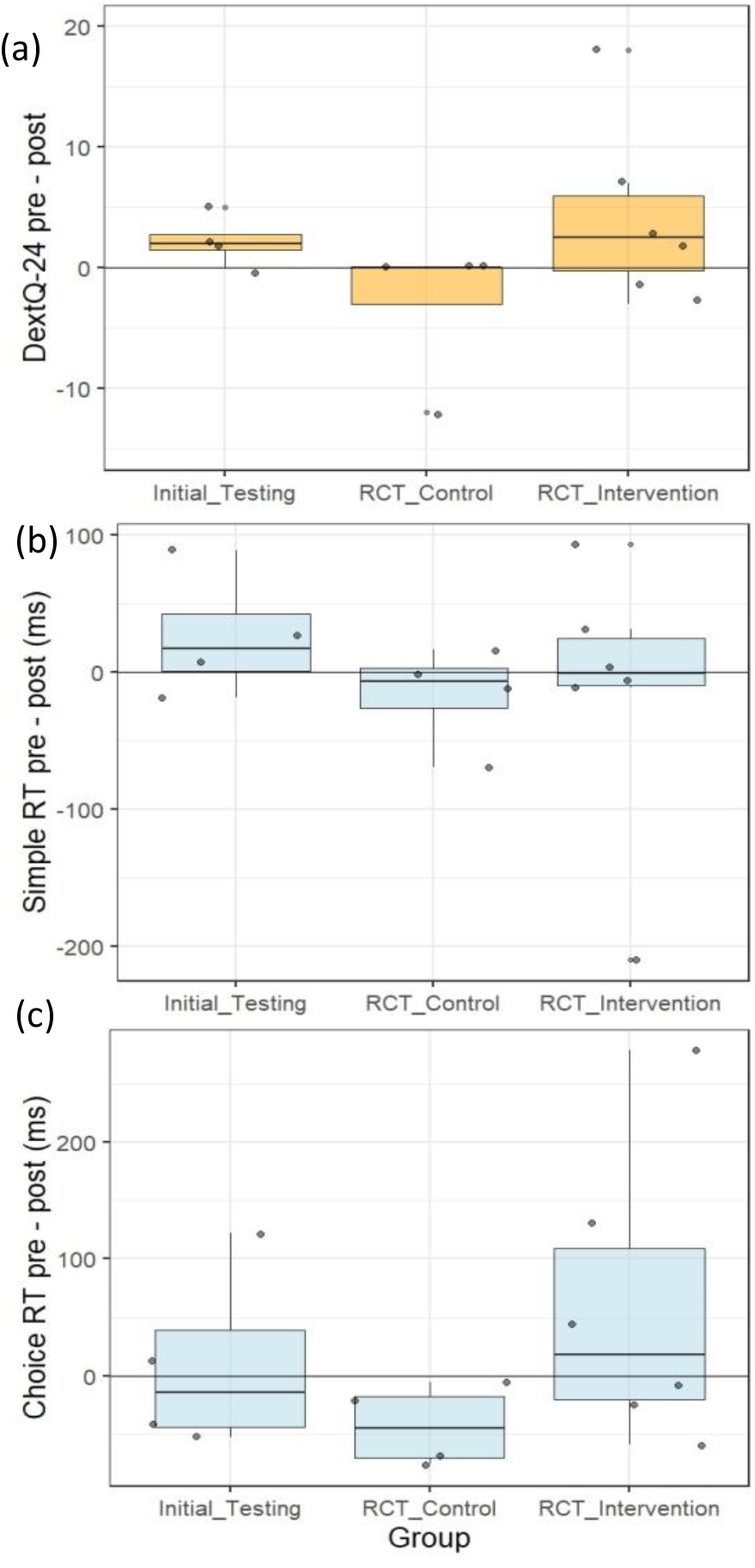
Median changes in exploratory outcome measures in the initial testing and pilot RCT: (a) DextQ-24; (b) simple reaction time; (c) choice reaction time. Boxes show quartiles and tails show 95% confidence intervals; dots represent individual participants. Positive change indicates improvement.

### Motor performance

Analysis of video-recorded action performance at baseline and post-intervention in the pilot RCT indicated reduced completion times for personally selected trained and untrained actions, and reduced difficulty ratings for all action types, in the intervention group (see Figure 6). In contrast, controls showed no evidence of improvement in completion times or difficulty ratings.

**Figure 6.**
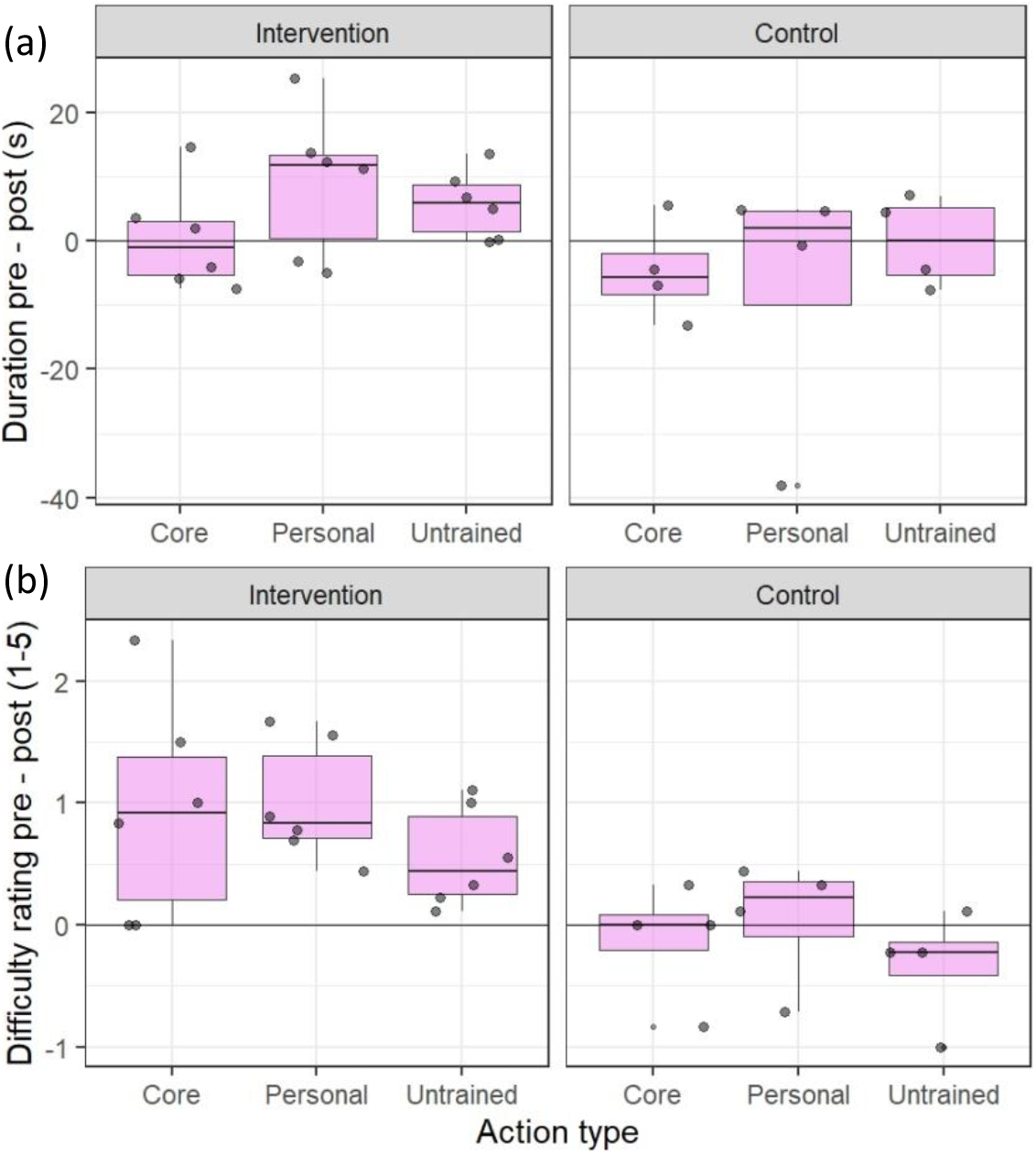
Median changes in (a) timed action performance and (b) difficulty ratings in the pilot RCT for the core actions (common across participants) and personally selected trained and untrained actions. Boxes show quartiles and tails show 95% confidence intervals; dots represent individual participants. Positive values indicate a post-intervention reduction in (a) duration or (b) difficulty.

## Discussion

ACTION-PD is a user-informed home-based intervention to improve everyday functional actions in people with PD, through combined action observation and motor imagery. The intervention was designed by an interdisciplinary team with input from people living with PD. A prototype app was developed to deliver the home training, with a simple interface intended to be user-friendly for people with PD, including those with limited experience of mobile technology. Given the heterogeneity and variability of PD, personalisation and flexibility are key elements of the intervention (see [7]). To obtain initial data on acceptability and usability, and to explore potential outcomes to include in a larger trial, we conducted a focus group and initial field testing, followed by a pilot randomised controlled trial. We note that the qualitative and quantitative findings described below are similar across both the initial testing and pilot RCT, despite some modifications to the intervention including the implementation of a new software platform.

### Acceptability and usability

The focus group indicated in-principle acceptability of the app and the proposed training protocol, while highlighting some potential barriers and limitations. In both phases of pilot testing, participants were able to use the app to train independently following initial set-up and guidance from the research team, as demonstrated for other home-based training programmes in PD (e.g., [10]). Initial testing indicated the need to slightly reduce the target training dose, which was subsequently achieved by all participants in the pilot RCT.

In addition to the usage data, the post-training interviews provided initial evidence that the ACTION-PD intervention is acceptable and usable for people with mild to moderate PD. Participants found the app and training protocol easy to use. The flexibility of the intervention allows individuals to fit the training into their daily routine and accommodate fluctuations in levels of fatigue or other symptoms, which participants appreciated. All participants expressed an interest in using a similar app in the future, and felt that the six-week duration of the current intervention was appropriate. While some participants found the actions well-suited to their needs, not all of the actions were considered to be sufficiently challenging. Indeed, it was suggested that the possibility of selecting new actions or moving on to more challenging actions could make training more motivating and sustainable. The focus group and post-training interviews also highlighted the value of feedback and encouragement to maintain motivation, consistent with previous findings in relation to other interventions for PD (e.g., [7,67,68]). Subjective ability to perform the motor imagery component of the intervention varied between participants. Some individuals found it difficult to engage initially but easier as training progressed, while others felt that their imagery did not improve over time. In this context, it should be noted that motor imagery ability varies widely among the general population [69], and although vividness of imagery is generally found to be preserved in PD, it may be affected in some cases [36].

Participants generally reported a preference for observing actions from the first-person perspective, although the overall contextual information provided by the third-person viewpoint was also appreciated. This corresponds with similar findings in stroke patients [51]. As noted above, people with PD may have difficulty in simulating actions from the first-person perspective [39,70]. The preference for the first-person video therefore suggests that, despite this potential difficulty, observation of an action from the first-person perspective may facilitate the generation of kinaesthetic imagery by providing a visual prompt, as highlighted in the following quote: “I’d feel more what that felt like to me, because the film was about…as if it was me that was doing the action”. This is consistent with the hypothesis that the role of AO in AO+MI is to provide an external visual guide for MI, as indicated by MI-specific effects on corticospinal excitability in healthy participants [71].

### Outcomes of AO+MI training in PD

Post-training interviews identified perceived improvements in performance of the trained actions, as well as other daily activities, indicating the potential to achieve broader benefits beyond task-specific training effects. However, some participants reported that improvements occurred early into the training period, with limited further progress, again highlighting the importance of progressive training.

Several participants reported using MI in everyday tasks following the training, such as dressing or getting out of bed. Additionally, the interviews indicated other ways in which AO+MI training may have influenced how participants approached actions. These included: (i) focusing attention so that tasks could be carried out in a more careful and controlled manner, as recommended in physiotherapy guidelines [72] and which speculatively could be linked to increased use of MI; (ii) reducing the stress associated with performing difficult actions; or (iii) highlighting subtleties of the movements. Potential psychological benefits including increased confidence and self-efficacy were also noted, consistent with other literature reporting these functions of motor imagery in older adults [73].

Analysis of action performance in the pilot RCT showed that completion times for both trained and untrained personally-selected actions were shorter following training in the intervention group, which corresponded to decreased difficulty ratings in the lab. This was broadly consistent with the pattern of difficulty ratings collected during training, which indicated that participants generally found the practiced actions easier by the end of the six-week period. However, most found the “core” actions selected by the research team less challenging than the “personal” actions that they had selected themselves, reinforcing the importance of personalisation.

Numerical trends in the exploratory outcome measures also suggested that the intervention could potentially produce improvements in dexterity and reaction times. We used a self-report measure of dexterity because of its direct relevance to the everyday actions targeted by the intervention, but in future trials this could be complemented by objective measurement tools such as a peg test [74]. A large-scale study of home-based dexterity training in PD [10] found improvements on both subjective and objective measures. However, to our knowledge, only one previous study has investigated effects of AO training - without MI - on dexterity in people with PD, where improved performance on a peg test was found [43].

Consistent with the findings from the interviews discussed above, in-app ratings of motor imagery in the pilot RCT did not show any subjective improvement across the six weeks. We also found no clear indication of improvement in motor imagery ability using the KVIQ across the cohorts. However, such self-report measures rely on the individual’s understanding of the concepts in question, and obtaining reliable pre/post data is dependent on consistent interpretation of the instructions over time. As noted above, qualitative data indicated that some individuals found it difficult to engage in imagery. Also, some participants showed an altered understanding of imagery as a result of the training, which may confound interpretation of scores on the KVIQ. Additional instruction and training in MI prior to the intervention might therefore improve understanding, engagement and consistency (e.g., [73]). Future work should also consider how best to evaluate effects of AO+MI training on the everyday use of motor imagery in people with PD, as commonly used tools assessing vividness of imagery (e.g., questionnaires such as the KVIQ) may not capture how imagery is used in the present intervention. Indeed, as noted above, some participants reported increased application of imagery to other everyday actions, beyond those practiced in training.

### Proposed mechanisms and future work

These preliminary findings demonstrate the potential for combined AO+MI training to facilitate everyday functional manual actions in people with PD. We can consider several mechanisms by which this may be achieved. First, specific motor representations for the trained actions may be developed or enhanced through AO and MI alongside physical practice (e.g., [75,76]). Second, the training may result in improved ability to generate MI for the practiced actions, such that participants are able to apply imagery more easily when performing the same actions outside of the training context. A third possibility is that participants develop stronger general skills in - or a greater awareness of - MI, which they are able to apply to functional actions beyond those practiced, as indicated by the improvement in timed and self-reported performance of untrained actions in the pilot RCT. Finally, as suggested by our qualitative findings, AO+MI training may lead to other changes in how actions are approached, such as focusing attention (e.g., [77]) or increasing confidence and self-efficacy (e.g., [73]). Indeed, combinations of the above may occur as a result of training, and these mechanisms should be further explored in future research.

Individual differences (for example, in motor imagery) may also influence the efficacy of home-based AO+MI training, such that some participants may obtain greater motor, cognitive or psychological benefits than others. In future, it may be appropriate to screen individuals to ensure a minimum level of MI ability prior to training, as in some previous studies of interventions for stroke (see [78]). Additionally, our qualitative data suggested that motivational factors vary between individuals, with some finding intrinsic motivation from the daily routine or the potential to improve their movements, while others may rely more on extrinsic motivators.

Themes relating to personalisation, variety and choice, and motivation, were echoed across the focus group and post-training interviews. In summary, key issues for further development of the intervention highlighted by the present findings include: (i) selecting appropriate actions at a suitable level of difficulty for the individual; (ii) offering variety, choice and progression in training; (iii) providing additional guidance or instruction to facilitate engagement in motor imagery; and (iv) increasing or maintaining motivation through the above as well as via positive reinforcement and feedback.

The present findings indicate that home-based AO+MI training delivered using mobile technology is feasible in people with mild to moderate PD. Based on the findings of this pilot work, a larger-scale randomised controlled trial should be conducted to evaluate the feasibility of the intervention, following further development with input from people with PD and healthcare professionals. Additionally, the involvement of healthcare professionals in selecting or prescribing appropriate training content, and delivering the intervention, should be considered. Our findings also have broader relevance for the development of behavioural interventions in PD, as well as applications of AO+MI in other groups, such as stroke survivors or healthy older adults.

## Acknowledgements

The authors would like to thank Elizabeth Barlow, Adam Lawrence, Kally McHugh-Simpson, Jade Pickering, and Nicole Rhodes for their assistance with data collection and coding, and Laurie Cooper and Digital Labs at Manchester Metropolitan University for software development. We also wish to thank all the participants involved in the focus groups and pilot testing.

