## Supplement S1 for "Action Imagery and Observation in Neurorehabilitation for Parkinson’s Disease (ACTION-PD): development and pilot randomised controlled trial of a user-informed home training intervention to improve everyday functional actions"

### Supplement S1: Themes from focus group

#### *Theme 1: Accessibility of technology*

Comments highlighted the need for application compatibility across a range of devices and operating systems, to ensure that the training is as widely available and accessible for people with PD as possible:

> *…I have a Windows phone, nobody makes apps for them. [FG2]*

Participants discussed how symptoms of PD could potentially interfere with the operation of the app, and that a simple interface would be appreciated:

> *I use an iPhone a lot, and fortunately, I've got very little tremor as yet, but I miss the keys, I misjudge it, very, very often, and I get the adjacent key to the one I want, and that's frustrating. [FG3]*

#### *Theme 2: Motivating influences: progression, encouragement and feedback*

It was noted that having a greater variety of actions to choose from would help to maintain motivation. In particular, the ability to progress from simpler actions to more complex sequences was suggested, and a simple game-like reward system could be incorporated:

> *I think to me, it's…about having progress. So that you get, you know, this is level one, and then you can always progress to level 2, if you've mastered that, as it were. [FG5]*

It was also acknowledged that rewards may be realised after longer-term use of the app, in that training may help to maintain the ability to perform actions that otherwise may diminish with disease progression:

> *With Parkinson's, you're actually working against the tide. So, in many ways, you need to be doing things before you lose the skills. It's about conserving, not just learning, isn't it. [FG5]*

The importance of positive reinforcement or encouragement was noted, which could take the form of either performance-based feedback or more simply recognition of regular practice.

> *We talked about the competitive element, didn't we, about knowing how well you were doing previously. We said that we'd like to know how we're doing. [FG4]*

> *Is there any sort of built in encouragement in, saying, particularly for people who do it every day, or, saying, oh great, you've done it, that's two days on the run. [FG6]*

#### *Theme 3: Personalisation and flexibility in training*

Participants discussed the importance of flexibility in the training protocol, such as in relation to the timing of medication, or to minimise fatigue effects:

> *A crucial feature would be the timing in relation to the last dose of tablets. [FG3]*

> *I'd like, for me, the idea of more small sections, more short sections, rather than a half hour block. Because it feels preferable to me, I could do a half hour block, but I think I'd start to be wandering by the end of it. [FG2]*

It was also noted that personal preferences might differ in terms of the device used to deliver the training:

> *I think the more flexible… it is, the better, because people will have different preferences. And because it's all about motivation, actually. [FG5]*

There was some discussion of features that could be incorporated within the app in future, such as reminders or music. It was acknowledged that these might be appreciated by some users but not others, and it was suggested that these elements could be optional:

> *You could have it like a permissive thing, if you could switch on a reminder. So if somebody who valued their reminder could do it, but it wasn't mandatory [FG5]*

> *I would want to be discreet, and not draw attention to the fact that I'm doing my exercises, so I don't want music playing [FG1]*
