## Supplement S2 for "Action Imagery and Observation in Neurorehabilitation for Parkinson’s Disease (ACTION-PD): development and pilot randomised controlled trial of a user-informed home training intervention to improve everyday functional actions"

Supplement S2: Everyday actions included in pilot testing

| Action name and description | Video duration (mm:ss) | Participants |
|---|---|---|
| Core actions | | |
| *Ticket sorting*: picking up a wallet from the table; opening the clasp; selecting and removing a train ticket; fastening the wallet; placing the ticket and wallet down on the table | | All Initial |
| *Coin sorting*: picking out two coins from a selection held in the palm of the non-dominant hand; transferring the coins to the palm of the dominant hand | | All Initial |
| *Water bottle*: unscrewing and removing the lid of a small water bottle; pouring water into a cup; replacing the lid; lifting the cup towards the mouth | 00:56 | All RCT |
| *Sugar jar*: unscrewing and removing the lid of a jar, extracting a teaspoon of sugar; depositing the sugar into a teacup; placing the spoon down; replacing the lid | 00:49 | All RCT |
| Personal actions | | |
| *Buttoning*: fastening and then unfastening 3 buttons on a tabard | 01:01 | All Initial; RCT3, RCT5 |
| *Food cutting*: using a knife and fork to cut 3 slices from a block of modelling clay shaped into a 'steak' | 00:50 | I1, I2 |
| *Butter container*: removing the lid of a plastic butter container | 00:16 | I2 |
| *Breakfast cereal*: opening a cereal box; pouring cereal flakes into a bowl; re-closing the box | 00:34 | I3 |
| *Lock and key*: inserting a key into a padlock; turning the key to unlock the padlock; *removing the key* | 00:34 | I4, RCT5 |
| *Coffee jar*: unscrewing the lid of a large coffee jar; screwing the lid back on | 00:48 | I4 |
| *Zipping*: fastening and then unfastening a zip on a jacket | 00:41 | RCT1, RCT5 |
| *Writing - patterns*: using a pen to draw pre-writing patterns on lined paper; maintaining the amplitude of the letters between the lines; completing 3 rows of different patterns | 00:52 | I1, I3 |
| *Writing- letters*: using a pen to write letters of the alphabet on lined paper; maintaining the amplitude of the letters between the lines; completing 3 rows of different patterns | 03:32 | RCT2, RCT3 |

| Action name and description | Video duration (mm:ss) | Participants |
|---|---|---|
| *Newspaper*: turning two pages or a newspaper; turning back to the cover | 00:45 | RCT1, RCT2 |
| *Shirt sleeves*: fastening 3 buttons on a shirt sleeve cuff; repeating on other sleeve | 00:54 | RCT2, RCT3, RCT4 |
| *Ticket sorting*: picking up a wallet from the table; opening the clasp; selecting and removing a train ticket; fastening the wallet; placing the ticket and wallet down on the table | 00:48 | RCT6 |
| *Ticket removing*: holding a wallet in one hand and set of tickets in the other; selecting a ticket; opening the wallet; placing the other tickets in the wallet; closing the wallet while holding the selected ticket | 00:44 | RCT6 |
| *Paper tidying*: gathering loose papers into a pile; picking up and straightening the papers; placing the pile back onto the table; picking up one sheet from the top | 00:48 | RCT6 |
| *Coin jar*: unscrewing the lid of a small jar and placing it onto the table; picking up 4 individual coins and dropping them into the jar; screwing the lid back on | 01:03 | RCT4 |
| *Yogurt pot*: removing the lid of a yogurt pot and placing it on the table; placing the lid back on | 00:37 | RCT1 |
| *Food bag*: opening a zip-lock food bag; removing a cookie and placing it onto a plate; re-sealing the bag | 00:44 | RCT4 |

*Note: I = initial testing cohort; RCT = pilot RCT intervention group.*
