## Supplement S3 for "Action Imagery and Observation in Neurorehabilitation for Parkinson’s Disease (ACTION-PD): development and pilot randomised controlled trial of a user-informed home training intervention to improve everyday functional actions"

### Supplement S3: Themes from post-training interviews

#### *Theme 1: Suitability and choice of actions*

Some participants noted that it was useful to practice everyday actions that would be commonly encountered. On the other hand, the similarity of the actions to the 'real-life' versions was questioned.

> *Well, I thought, they were probably spot on really cause they, sort of, hit the problems that people do have, you know, the fastening of buttons and zips and very common place problems [I5]*
>
> *…the worst is if I'm out and about and there's more pressure on me and I'm there in front of the ticket collector and he's got a big queue gathering behind me and I'm … slow, and that's where things really go pear shaped [I6]*

Several participants reported that the actions were unchallenging, or that they found only one or two of the actions difficult. Other participants found the actions better suited to their needs, or appreciated the combination of easier and more difficult actions.

> *I think I regretted that I hadn't chosen some of the other actions on the list that was given to me, which might have been a bit more challenging so that I could have practiced those and maybe noticed an improvement in those or not, as the case may be, and I think I picked ones that were too easy really [I2]*
>
> *I found some easier than others but that didn't really matter and, in a way, I used to sort of shuffle the order of what I did quite a lot, the easier tasks almost gave me a bit of a pause [I6]*

*Nonetheless,* all participants felt that the intervention would benefit from having a greater variety and choice of actions, and it was suggested that participants could be supported to select actions appropriate to them. Participants would like to have the option to replace actions once a level of competence had been achieved, or to be able to progress to more difficult actions.

> *You or the person setting up the app, would probably need to, somehow identify with my help, which actions I can really benefit from, before embarking on the six-week test……make them progressively more difficult, maybe. You could have the writing going on to the end of the alphabet or extend it in some way [I2]*
>
> *It needs to have a larger variety of different activities, videos...You need like a menu of all the different actions and then you just pick out half a dozen. Then, when you feel you've mastered those, you can pick out some more. (P3)*

One participant felt that they would prefer to focus on one action at a time, according to their current needs.

> *I might have 30 different actions on the thing and I'd think, hmm, this is one I've been struggling with recently, and I'd just work on that one for two weeks. And then, yeah, I could select another if I needed it and work through that and then maybe check back every so often to see if the skills are still there [I6]*

#### *Theme 2: Action observation and motor imagery*

It was noted that watching the videos provided useful cues for improving performance, and one participant noted that the videos were particularly helpful for the more difficult actions. It was also suggested that watching the videos could increase awareness of variability in the observer's own actions. However, one participant noted that they became distracted while watching the videos.

*…the films help you get into the right mind-set as well as watching again how to do it well or just how to do it in an easier manner…I think what helped was doing it so often you noticed little things which could make a difference potentially. (P4)*

*…watching the way the she did it and the way I did it, I know that the way I did it varied a lot. (P2)*

Participants generally reported a preference for the first-person perspective, but this could change over time. Comments indicated that the first-person video may have promoted motor imagery, although some participants appreciated seeing the third-person view first to get an overall understanding of the action. Some participants felt that it was helpful to see both perspectives, which might facilitate motor imagery and learning.

*When I first saw them, the first one, it's like the third person. I thought that was fine. Then, when I saw it in the first person, I thought that's better. But, after a while, I just thought, no, there's no difference between them really. (P3)*

*the views of the first person they were more helpful than the view facing the person but maybe that was necessary to give context to the exercise anyway [I6]*

*It was useful to have the two different perspectives and that helped with the visualising of the thing. [I2]*

*To help you re-learn tasks, having it from the two angles would be better. [I3]*

Participants reported mixed experiences of the motor imagery component of the training, but this may have improved over time.

*At first, I was really concentrating on what had to be done but as time went on, as it became easier, there was less thought going into it…it became a natural part of it [I4]*

Some participants found it effortful to engage in motor imagery, while other comments indicated that the importance of the imagery component might be unclear.

*I put it down to practice and really trying, trying very hard. Sometimes, just closing my eyes and trying to imagine the action or the feeling. (P3)*

*I'm not sure how much I was really accessing the imagery or whether I was just really playing the game as it were, really. It was hard to say how much I was really deriving from this concept imagery [I2]*

Hearing the sounds associated with the actions may have helped to facilitate imagery:

*…sound effects were definitely helpful, I think without the sound there was a bit more distance between myself and the image, with the sound it was more as though I was there [I6]*

*Theme 3: Accommodating the training into everyday life with Parkinson's*

*Subtheme 3.1. Acceptability of time demands*

Participants generally found the training schedule manageable, and felt that they were able to accommodate the practice into their day, valuing the flexibility of the intervention.

*I think in terms of the number of exercises, that was about right. If you made it six or seven a session, I think that would be too much….at times, it was quite hard to fit in one in a day. But then there were other…I mean, I did two in a day in some cases. (P2)*

*I liked the fact that you could do it whenever you had time to do it. For me on some weeks that was at different times of the day, and I think if it had been rigid it would have been more difficult to fit it in (P4)*

However, one participant commented on the additional time needed to set up the objects in preparation for their session:

*Even though it only takes about 15 minutes, all your preparation and then putting it all there afterwards and that, takes up a good half hour, I found, really. …. when you're busy, you've got to find the space and the time to do it in… I enjoyed doing it, but I didn't enjoy doing the preparation for it. (P3)*

Another found the schedule more demanding and found that they had sacrificed other activities in order to fit in their training:

*It did take quite a chunk out of the day, to get it in I dropped off the physio exercises which probably...I mean I do those about three times a week, that's probably not a good thing to drop off [I6]*

The duration of the current intervention was found to be manageable and appropriate:

*I think six weeks was enough, otherwise I think it would get too, how can I put it? A chore, rather than…I don't know what word to use, yes, quite a chore rather than…you wouldn't enjoy it so much [I4]*

*Subtheme 3.2. The impact of medications or symptoms on training.*

Some participants noticed that their performance was affected by medication effects or fatigue.

*And I tried doing those when I was…like at different times and when I knew I was short of medication, and that did take a long time. (P2)*

*my hand movements suffer when I'm not fully charged with medication and I did find that the actions that I had to do were sometimes a bit more difficult when I was off my medication [I2]*

*I found that if I left it late in the day, it was more tiring than doing it in the morning, as in my energy levels would run out for the day [I3]*

The variable nature of Parkinson's and the way it can affect different actions was also commented on by several participants:

*I did find it more difficult if I'd had dyskinesia, it was very difficult to do things like the zip …if you're having a bad day, you might have done very well the day before, say, on the zips and the next day you might be absolutely rubbish, so you don't get a consistency, I don't think, you get good days and bad days [I5]*

<u>*Theme 4: Perceived effects including cognitive and psychological changes*</u>

*Subtheme 4.1: Performance, mindset and imagery.*

Most participants noticed at least some improvement on the actions trained within the intervention, although some did not experience any improvement, which possibly related to the suitability of the selected actions.

*From my point of view that there's not much doubt it had a beneficial effect [I6]*

*I felt they're split almost into two categories, I had two that I found very straightforward and simple and I had three that were quite difficult. They never seemed…you know, none of those really ever changed at all (P1)*

The training had helped some participants in performing other everyday tasks. Comments suggested a change in approach or mindset when performing actions:

> *It made me a bit more attentive about the everyday things, and I wasn't expecting to have that benefit…Even things like I struggle changing gear and I just thought well, maybe you're putting your hand in the wrong position, maybe in a different position it might be helpful…opening a bottle of milk or opening the door or any of those things that the actions related to it made things a bit more helpful in everyday life. (P4)*

> *…putting the mind into action before I do the task …it doesn't take that long to…it's not a deep thought, it's just a process of what I've got to do [I4]*

Some comments more explicitly referred to changes in awareness or use of action representation processes (observation and imagery) in everyday life, although some participants did not notice any such changes. Specific examples included applying imagery to tasks including tool use, dressing, getting out of bed and moving through doorways.

> *The idea of imagery hadn't entered my knowledge, sort of thing. And with the way of imagining things happening, I think that's given me an insight into other things, so as a spin-off from it…I think possibly subconsciously you're analysing what you're doing in all the tasks as if you've been watching a video [I3]*

> *when I'm getting dressed one thing I always find quite difficult is putting a leg into trousers... I try to imagine what I'm doing and then remove the leg before I tried to actually do the full action [I1]*

*Subtheme 4.2:* Psychological effects

Other changes such as increased confidence and sense of control were reported by some participants.

> *…doing this helped me realise that there's some things I could work on and potentially then control sometimes how I do something. So like a simple action like turning the coffee jar, originally I thought I just can't do that anymore, or it's just difficult to do it, whereas practicing it made me think actually you can control things a little bit more than you think you can and not accept that you think you can't do something (P4)*

> *I found that practicing took away an element of the stress and made the whole thing a bit easier and a bit simpler to do. So, I found it quite helpful, in that regard [I2]*

<u>*Theme 5: The importance of motivation and feedback*</u>

Motivation was unanimously considered an important issue, although participants' views on what would motivate them differed.

For some participants the potential to improve movements through the training, or just completing the daily sessions, was intrinsically motivating.

> *The motivation is learning how to do something that you've forgotten how to do. I think that's motivation in itself [I3]*

> *Well every day it was quite an achievement really, for me, doing things on a daily basis once again [I4]*

Practicing more challenging actions, or a progression in the difficulty of actions might also provide a source of motivation:

*…once you'd done those actions and you got confident you could do them very competently…to go on to something more challenging, different or challenging [I1]*

Some participants would find performance-related feedback helpful. It was also suggested that more feedback and encouragement could be built into the app:

*It comes back to the old school days, if you're doing a thing, you like to know what progression you've made and if you're not doing it, it would be nice to find out what needs to be done to improve [I4]*

*I'd like a bit more…words on the screen, telling you how you're doing and what to do next and then encouraging you, with the option of a voice-over, doing the same thing… Depending on which answer you tap, it could then say, well don't worry, keep going at it. Or, it could say, well done, that's good. (P3)*
