## Supplement S4 for "Action Imagery and Observation in Neurorehabilitation for Parkinson’s Disease (ACTION-PD): development and pilot randomised controlled trial of a user-informed home training intervention to improve everyday functional actions"

Supplement S4: Difficulty ratings and motor imagery ratings during home training

(i)

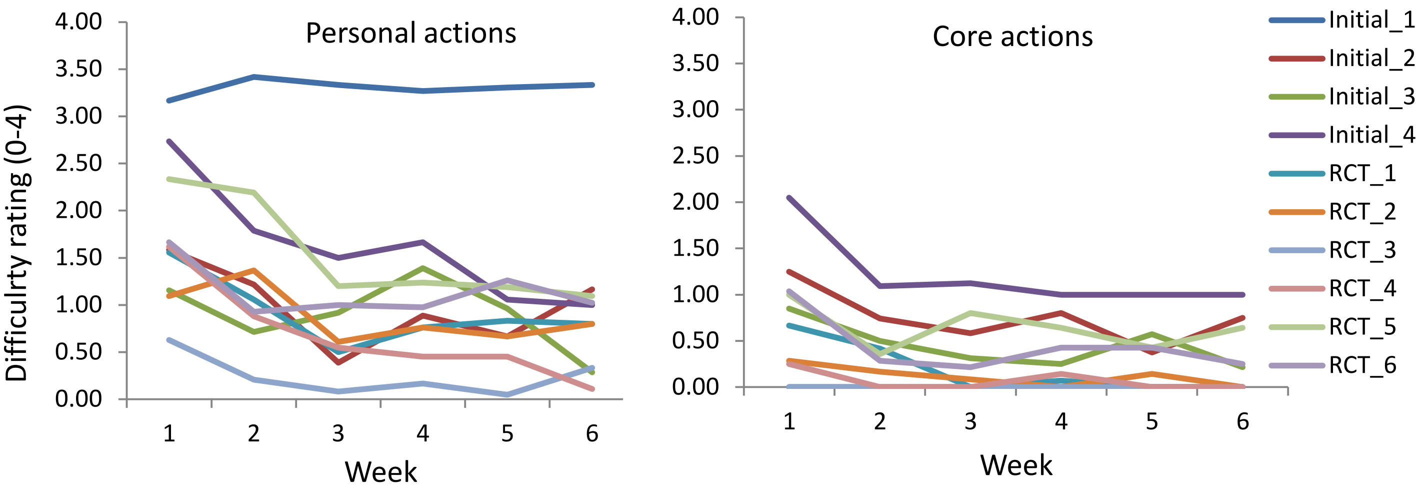


(ii)

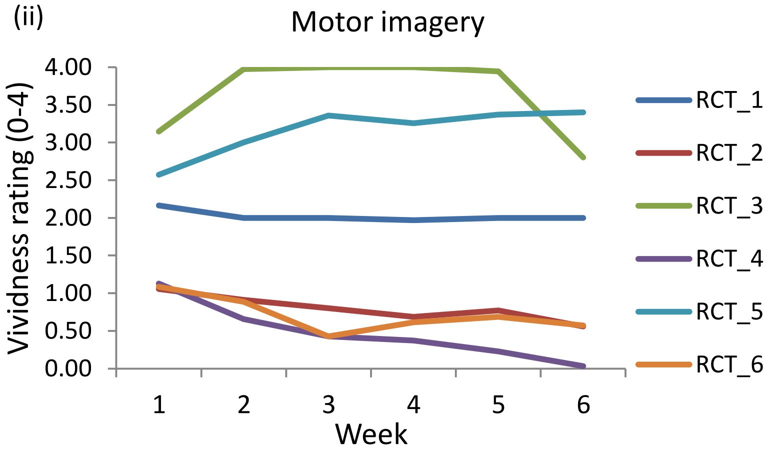


*Figure S4.* Average weekly ratings during the six-week intervention for (i) difficulty of personal and core actions in the initial testing and pilot RCT; (ii) motor imagery in the pilot RCT.
